# Light-induced Extracellular Vesicle Adsorption

**DOI:** 10.1101/2024.04.24.590318

**Authors:** Colin L. Hisey, Xilal Y. Rima, Jacob Doon-Ralls, Chiranth K. Nagaraj, Sophia Mayone, Kim T. Nguyen, Sydney Wiggins, Kalpana D.P. Dorayappan, Karuppaiyah Selvendiran, David Wood, Chunyu Hu, Divya Patel, Andre Palmer, Derek Hansford, Eduardo Reategui

## Abstract

The role of extracellular vesicles (EVs) in human health and disease has garnered considerable attention over the past two decades. However, while several types of EVs are known to interact dynamically with the extracellular matrix and there is great potential value in producing high-fidelity EV micropatterns, there are currently no label-free, high-resolution, and tunable platform technologies with this capability. We introduce Light-induced Extracellular Vesicle Adsorption (LEVA) as a powerful solution to rapidly advance the study of matrix- and surface-bound EVs and other particles. The versatility of LEVA is demonstrated using commercial GFP-EV standards, EVs from glioblastoma bioreactors, and E. coli outer membrane vesicles (OMVs), with the resulting patterns used for single EV characterization, single cell migration on migrasome-mimetic trails, and OMV-mediated neutrophil swarming. LEVA will enable rapid advancements in the study of matrix- and surface-bound EVs and other particles, and should encourage researchers from many disciplines to create novel diagnostic, biomimetic, immunoengineering, and therapeutic screening assays.

---

Extracellular vesicles (EVs) are micro- and nanoscale lipid-enclosed packages that are released by all cells and are found in various biofluids and tissues[1–3]. Due to their ability to relay bioactive cargo over short and long distances or deliver therapeutic payloads, they have experienced nearly two decades of increasing study aiming to better understand their dynamic roles in health and disease[4, 5]. EV research has grown rapidly and benefited greatly from significant efforts to enhance rigor and reproducibility[6]. However, most studies have focused on EVs isolated from biofluids or conditioned media of cell cultures rather than EVs present in or on the extracellular matrix (ECM). To fully explain the role of these understudied EVs and enable the creation of novel EV characterization assays, a nonspecific, tunable, rapid, and high-resolution method of EV micropatterning is needed[7].

To date, EV micropatterning has largely been developed for use in single EV characterization assays as an alternative to ensemble techniques, primarily for diagnostic applications. These studies often combine complementary characterization tools to classify EVs based on their surface antigens, internal contents, or even physical and mechanical properties[8–10]. Of these techniques, single EV characterization using fluorescence requires efficient EV capture using antibodies, other specific ligands, or nonspecific lipoplexes to effectively concentrate EVs on surfaces[11–13]. Then, total fluorescence or colo-alization of multiple fluorescent probes (e.g. antibodies, molecular beacons) can be used to discover latent patterns within different EV populations[14–17]. To avoid potential bias from selective capture approaches[18], a tunable and nonspecific approach to EV micropatterning could pave the way for novel single EV characterization assays, including controlled titration of EV surface density or engineered mimicry of ECM-bound EVs[19].

Several key stimuli within microenvironments are known to influence cell behavior, such as chemical gradients, fluid flow, mechanical properties, and topographical cues[20–22]. EVs within the extracellular matrix (ECM) have also been shown to contribute to these dynamic stimuli and mediate inter-cellular communication, playing uncertain roles in processes ranging from metastasis to angiogenesis and wound healing[23–27]. A more complete understanding of these EVs could lead to the development of novel therapies and improved clarity of their function. For example, one recently discovered and understudied type of EV, the migrasome, is left behind during migration in “breadcrumb” trails in a process known as migracytosis. Deposited migrasomes can then be internalized by trailing cells[28–30] and they have even been found in circulation, indicating that they must detach from the ECM with some regularity[31]. However, their relationship to classical matrix-bound EVs and other tissue-derived EVs remains unclear[32], and this ability for ECM-bound EVs to act as cellular messengers is known solely by observation as there are no versatile and scalable methods for precisely micropatterning migrasomes or other ECM-bound EVs in vitro.

Additionally, EVs influence the dynamic responses of immune cells by mediating antigen presentation [33, 34], suppression [35, 36], and immunity [37–39]. Neutrophils are highly abundant circulating leukocytes that actively engage and destroy pathogens via chemotaxis, adherence, phagocytosis, and NETosis[40, 41]. At sites of infection or injury, neutrophils are the first responders, where an initial neutrophil will recognize this site and begin to recruit additional neutrophils and initiate an immune signaling cascade. This event of coordinated migration is called neutrophil swarming[42, 43]. To better understand this process, several in vitro platforms have been developed to micropattern antigen-functionalized particles or whole cells and then quantify the swarming behavior of added neutrophils[44–47]. However, despite evidence that bacterial EVs, or outer membrane vesicles (OMVs), serve diverse roles in immunity ranging from probiotic to pathogenic[48–51], no studies to date have reported neutrophil swarming mediated solely by OMVs. Thus, a technology that can precisely micropattern OMVs onto conventional cell culture surfaces could be used as a platform for developing tunable immunoengineering assays involving EVs and other particles[52].

To address these and many other technical challenges in the EV field, this study details the development of a versatile platform technology called Light-induced Extracellular Vesicle Adsorption (LEVA). LEVA enables the high-resolution, nonspecific, tunable, scalable, and rapid micropatterning of EVs from diverse sources for a wide range of applications in human health. Initially, LEVA was optimized using large EVs (lEVs) and small EVs (sEVs) from bioreactor cultures, as well as commercially available GFP-EV standards, to produce well-defined shapes and gradient patterns at subcellular resolution. COMSOL simulations and time-lapse total internal reflection fluorescence microscopy (TIRFM) imaging were then used to quantify the underlying EV adsorption kinetics and its dependence on EV size. Next, LEVA was applied to several novel EV applications including the first reported examples of digitally titrated single EV characterization by protein and RNA fluorescence colocalization, patterning of migrasome-mimetic trails followed by single cell migration, and E. coli OMV-mediated neutrophil swarming. As LEVA exploits relative increases in attractive forces between the engineered regions of surfaces and innate EV properties, it fundamentally relies on nonspecific adsorption rather antibody or other specific ligand-based capture. Thus, it can likely be applied at scale to many other types of applications with little modification to the reported workflow. Finally, such precise and tunable control of the location and concentration of EVs on surfaces should ultimately enhance rigor and reproducibility within this rapidly growing field.

## 1 Results

### 1.1 Light-induced Extracellular Vesicle Adsorption

Based on the initial template design, the DMD photochemistry process exposes certain regions of the PLL, mPEG-SVA, and PLPP-coated surface to UV illumination for durations that scale linearly with the template’s grayscale values at nearly single micron resolution (detailed in Methods section). This photocleavage results in well-defined regions on the surface with predictable levels of nonspecific EV adsorption. A representative schematic of a typical experimental workflow is presented in Fig 1A, including an optional iterative approach for patterning multiple types of EVs. An example of a grayscale template and the resulting EV micropattern following the LEVA process are shown in Fig 1B and C, respectively. In these templates, black regions represent areas which are not exposed to UV and will fully retain PEG at their surface, whereas white/gray regions are exposed for a duration that scales with their grayscale values. An additional image of EVs patterned as an EV crosssection image, with magnified regions showing nearly single EV resolution patterning are shown in Fig 1D, and an example of multi-EV patterning is shown in Fig 1E.

**Fig. 1.**
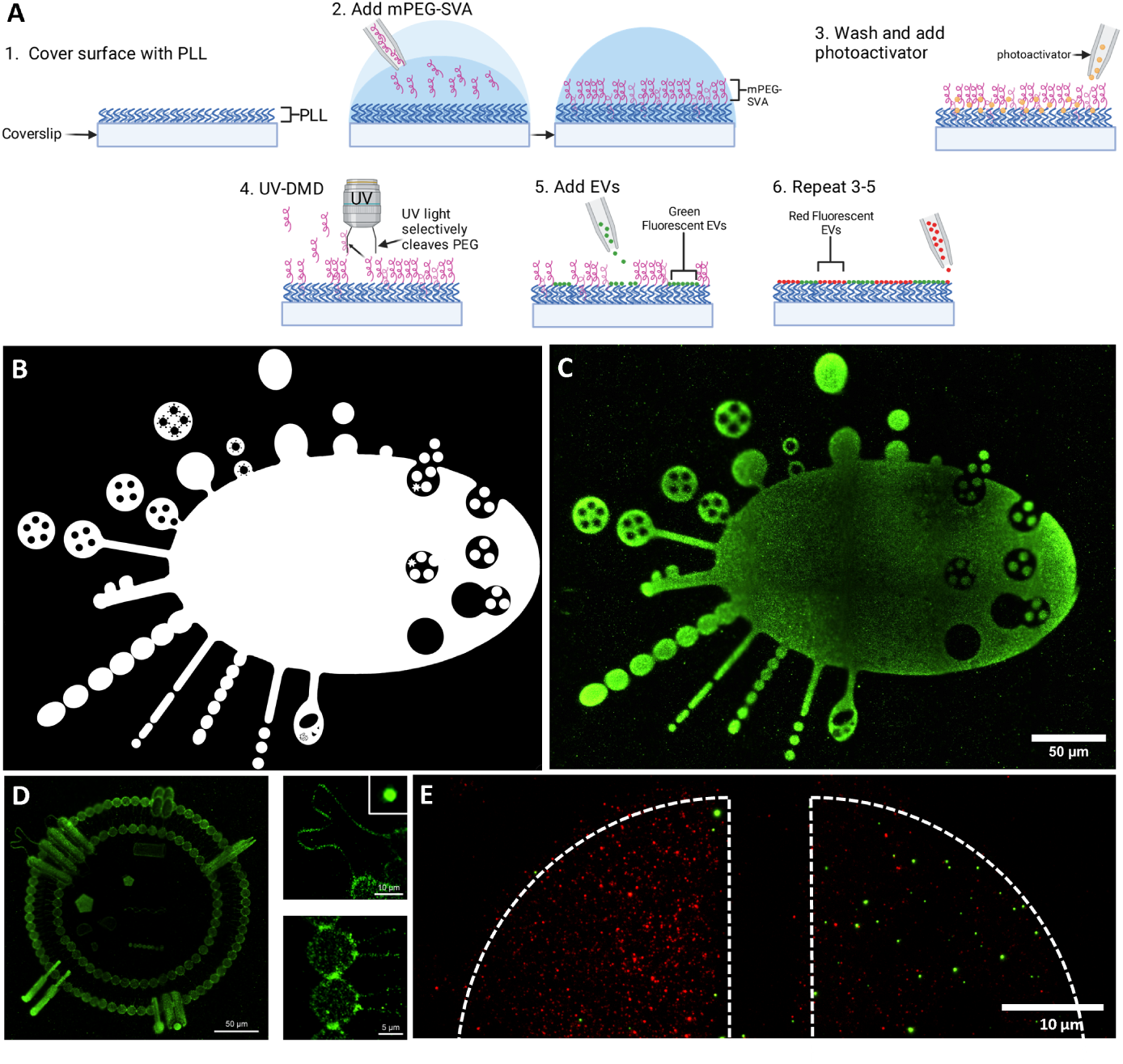
Light-induced Extracellular Vesicle Adsorption (LEVA). (A) LEVA schematic showing coating of an oxygen plasma-cleaned coverslip with PLL, mPEG-SVA, and a high concentration photoactivator. (B) Example of patterning template image adapted from Buzas et al.[52]) and (C) resulting EV micropattern using WGA-labeled U-87 MG sEVs (scale bar = 50 *µ*m). (D) U-87 MG sEVs patterned as an EV crosssection with magnified regions showing nearly single EV resolution patterning (image adapted from Almeria et al. [53]) and (E) an example of multi-EV patterning showing the initial EV pattern on the left (red) and the second EV pattern on the right (green).

The stark contrast in binding affinity between these different regions, combined with the ability to rapidly produce gradients by dynamically modulating UV exposure, ultimately enables the selective adsorption of EVs with high predictability and customization. Furthermore, the PEG-functionalized regions greatly reduce initial nonspecific binding of EVs and allow rapid removal of most of the undesired EVs with simple wash steps without significant loss of EVs in the desired regions. Importantly, by not employing antibodies or other ligands for EV capture, the platform greatly reduces cost and increases scalability compared to any previously reported technologies.

### 1.2 LEVA Kinetics and Simulations

To better understand the LEVA process, COMSOL Multiphysics simulations and time-lapse TIRFM were used to model and experimentally measure EV binding kinetics on the engineered surfaces. Fig 2A and B show the size distributions of U-87 MG bioreactor-derived lEVs and sEVs, respectively, which were characterized using nanoparticle tracking analysis (NTA) and negative staining TEM to verify their expected size and morphology[6]. It is important to note that the optimization and development of this process was largely made possible by first producing vast quantities of EVs using CELLine AD 1000 bioreactors and treating EVs as an expendable material[54, 55].

**Fig. 2.**
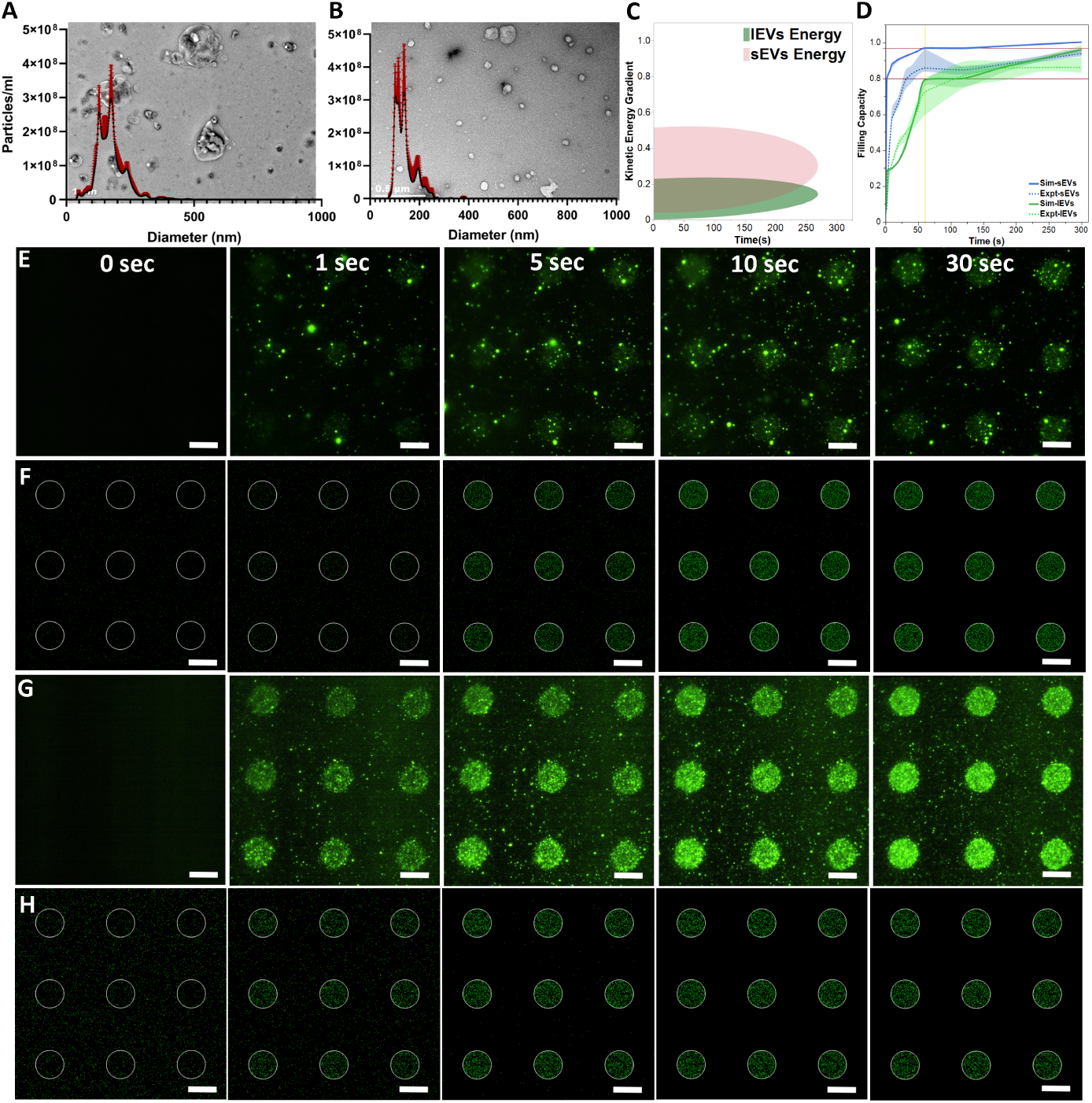
Study of large and small EV binding kinetics in LEVA. (A-B) NTA size distribution and TEM images of bioreactor-produced U-87 MG lEVs and sEVs. (C) Kinetic energy and (D) particle density over time based on COMSOL Multiphysics particle simulations lEV and sEV-sized particles. Time-lapse TIRFM microscopy images and COMSOL Multiphysics simulations results of LEVA for (E-F) lEVs and (G-H) sEVs (scale bar = 10 *µ*m).

Simulations were performed based on estimated properties of these two general EV subpopulations, revealing an exponential increase in particle density at the positively charged surface regions, while density decreased over the rest of the glass surface where the charge remained neutral. Notably, sEVs (mean diameter = 100 nm, standard deviation = 50 nm) exhibited a higher rate of particle binding capability compared to lEVs (mean diameter = 135 nm, standard deviation = 45 nm). This was attributed mostly to the increased effect of Brownian motion on sEVs compared to lEVs, resulting in greater mobility and a higher probability of contacting the patterned regions on the surface. sEVs took approximately 8 sec to sufficiently bind and settle on the positively charged surface, whereas larger EVs required around 58 sec. Increasing the average size of the EVs resulted in even longer binding times. Smaller EVs also bound more closely to the positively charged surface, allowing for a clear delineation of the surface boundary, while larger EVs exhibited slower binding velocities, impacting their kinetic energy. Fig 2C and D show the kinetic energy and particle density over time for both lEVs and sEVs.

To validate the simulation results experimentally, bioreactor-produced U-87 MG lEVs and sEVs were labeled with WGA-488, free dye was removed by repeated ultrafiltration and PBS washing, EVs were normalized by concentration using NTA, and EVs were added instantaneously to the patterned arrays of 10 *µ*m circles in a manner consistent with the simulations. Time-lapse TIRFM was used to monitor the EV adsorption in real time. Remarkably, the quantified fluorescence intensity results from TIRFM videos agree with the simulation results, showing slower binding kinetics of lEVs (Fig 2 E-F) compared to sEVs (Fig 2 G-H) shortly after the addition of the EV solutions. As illustrated in Fig 2, while the rapid TIRFM acquisition of images resulted in some photobleaching of the EVs, there is clear and rapid initial adsorption of EVs in the UV-illuminated, or positively charged, regions with minimal nonspecific binding to the unexposed areas. Videos of COMSOL Multiphysics simulations and time-lapse TIFRM used for both lEV and sEV analyses can be found in Supplementary Videos 1-4.

### 1.3 Resolution Test

To determine the resolution limits of LEVA, several types of fluorescently labeled EVs were patterned using identical templates and workflow. To ensure that the reported results are from EV binding and not simply dye contamination, commercially available GFP-EV standards were also used. Additionally, a comparison between WGA-labeled U-87 MG sEVs and a WGA-only control are shown in Supplementary Figure 1. As with any fluorescent labeling of EVs, there is a slight background fluorescence with the dye-only control, but it is negligible compared to the sEV sample. The initial templates for resolution testing included 1, 2, 5, and 10 *µ*m diameter circles as shown in Fig 3A and 1, 2, 5, and 10 *µ*m microtracks as shown in Fig 3B. Generally, consistent EV patterning results were only seen for 5, and 10 *µ*m diameter circles, while 2, 5, and 10 *µ*m microtracks were consistently patterned successfully. EVs could also be patterned on 2 *µ*m circles on occasion, but 1 *µ*m circles and microtracks were rarely visible despite repeated attempts. TIRFM images of successful EV patterning are shown Fig 3 for (C) U-87 MG sEVs, (D) U-87 MG lEVs, (E) E. coli OMVs, and commercially available GFP-EV standards from (F) Vesiculab and (G) Sigma. Clearly, depending on the EVs used, the fidelity of the EV micropatterns can vary, but successful EV micropatterns were produced for all the samples tested in the desired regions, with minimal nonspecific binding in the unexposed regions.

**Fig. 3.**
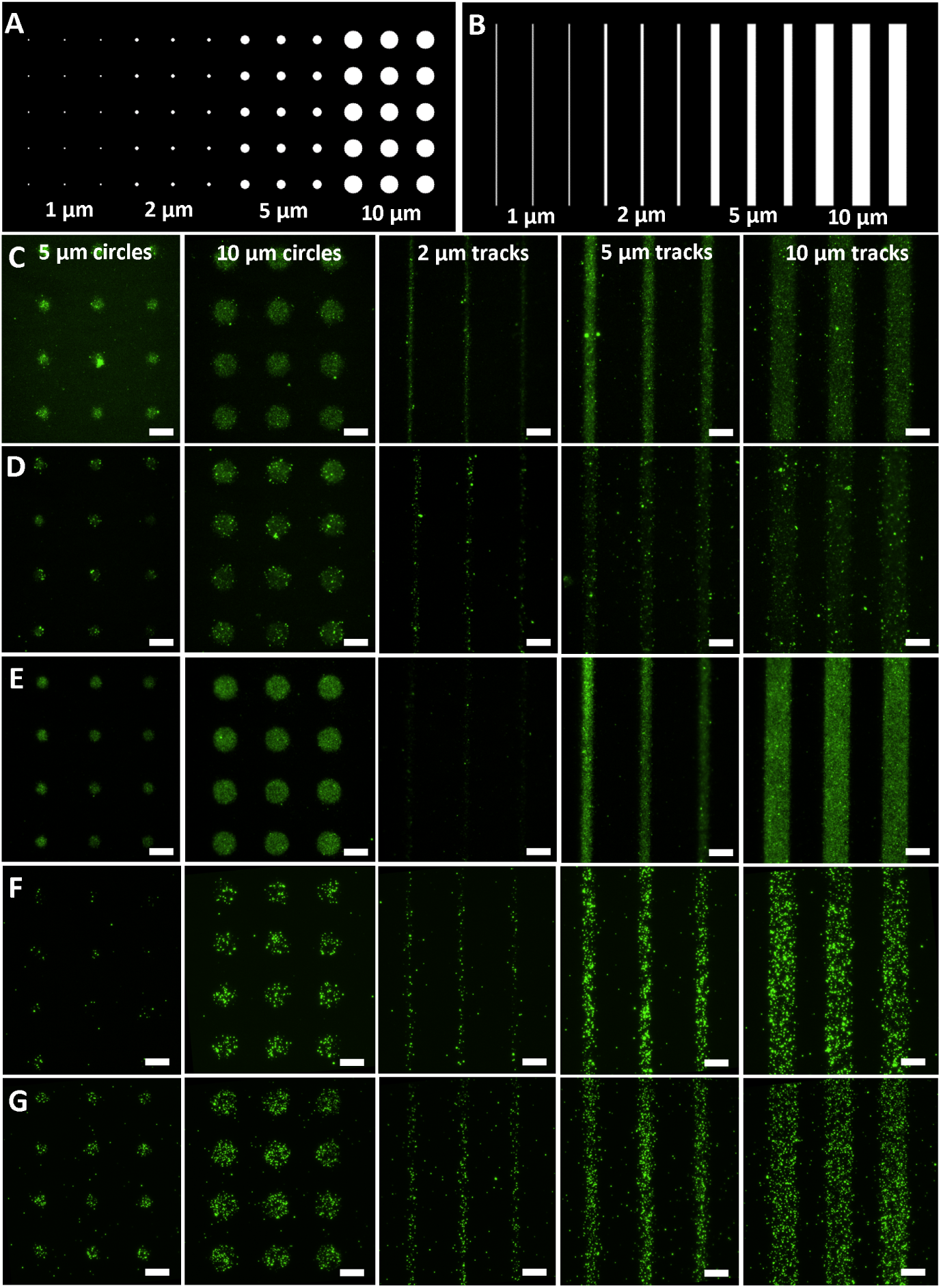
LEVA resolution test using various EV sources. (A) Resolution test template for circle array. (B) Resolution test template for microtrack array. TIRFM images of successful LEVA micropatterns for (C) U-87 MG sEVs, (D) U-87 lEVs, (E) E. coli OMVs, and commercially available GFP-EV standards from (F) Vesiculab and (G) Sigma. (scale bars = 10 *µ*m)

### 1.4 EV Digital Titration

We attempted to use gradient templates to micropattern EVs from various sources. Three gradients were chosen to determine the adsorption differences, including a linear gradient to visualize chromatic increases in free energy, an exponential decay gradient to visualize drastic changes in gradation, and a Gaussian distribution to simulate the stochastic accumulation of particles towards a central line. Generally, the sEVs produced more homogeneous patterns, whereas lEVs produced more variable patterns with higher rates of non-specificity as shown in Fig 4. For the exponential decay gradient shown in Fig 4A, the sEVs demonstrated a better fit (Spearman’s coefficient, p = 0.9983 for sEVs and p = 0.9749 for lEVs, p *<* 0.0001) and for the linear gradient shown in Fig 4B, sEVs again demonstrated the closest fit (Spearman’s coefficient, p = 0.9998 for sEVs and p = 0.9943 for lEVs, p *<* 0.0001). Finally, for the Gaussian distribution shown in Fig 4C, both sEVs and lEVs had similar fits, with sEVs yielding a slightly better fit (Spearman’s coefficient, p = 0.9942 for sEVs and p = 0.9926 for lEVs, p *<* 0.0001). Interestingly, at lower points of the gradients, the lEVs consistently demonstrated greater adsorption Fig 4. Although the EVs did not perfectly reproduce the patterns, with the edges of the pattern demonstrating decreases in fluorescence, the EVs followed the desired trend and demonstrated spatial control not currently achieved in any reported techniques. Quantification of the relative mean fluorescence intensity for the U-87 MG lEVs and sEVs compared to the desired template are reported in Fig 4D, indicating that in the future, the templates can be tuned for specific types of EVs in order to produce the intended pattern more precisely.

**Fig. 4.**
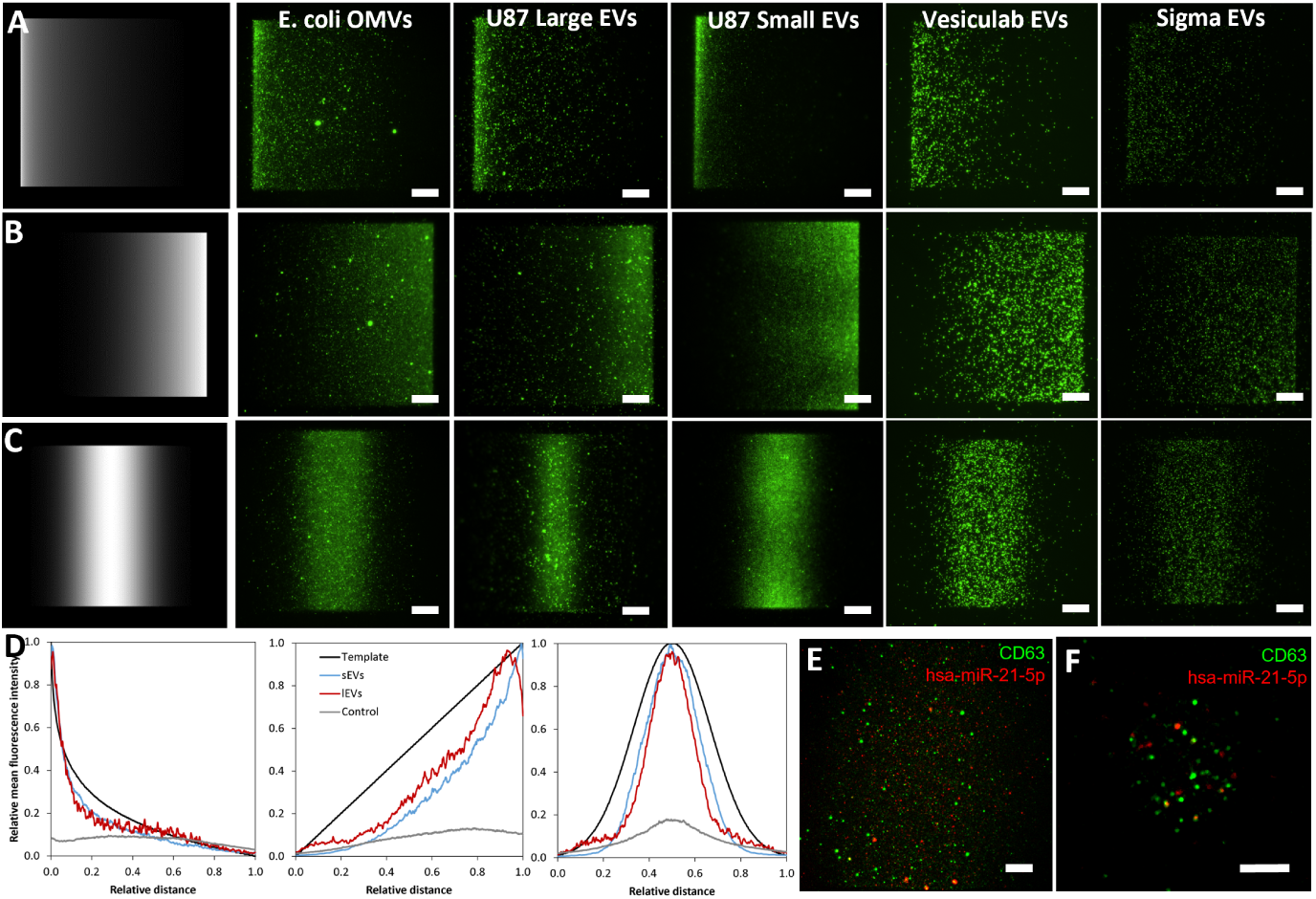
LEVA gradient test using various EV sources. (A) (left to right) Exponential gradient template and resulting patterns for E. coli OMVs, U-87 MG lEVs and sEVs, and commercially available GFP-EVs from Vesiculab and Sigma. (B-C) Linear and Gaussian gradients templates and resulting TIRFM images in a similar manner (scale bars = 15 *µ*m). (D) Quantification of desired EV patterning density and normalized fluorescence from TIRFM images for exponential, linear, and Gaussian gradients for U-87 MG lEVs and sEVs. (E) Digital titration of anti-CD63-AF488- and miR-21 molecular beacon co-labeled U-87 MG sEVs (scale bar = 15 *µ*m) and (F) similarly colabeled U-87 MG lEVs but in a in a 10 *µ*m circle pattern (scale bar = 5 *µ*m).

As a proof-of-concept, U-87 MG EVs were also co-labeled with anti-CD63-AF488 antibody and hsa-miR-23-5p molecular beacons and patterned to determine whether LEVA could be used in future colocalization studies. Unlike conventional ELISA assays, wherein antibodies or other ligands are used to first capture EVs of interest, and fluorescent probes are then added to label the EVs while blocking nonspecific binding, EV digital titration requires pre-labeling EVs in solution followed by a purification step to avoid deposition of unbound probes that are indistinguishable from the EVs. An example of successfully pre-labeled and micropatterned U-87 MG sEVs in the Gaussian distribution are shown in Fig 4E, and U-87 MG lEVs micropatterned in a 10 *µ*m circle are shown in a Fig 4F.

### 1.5 Migrasome-mimetic EV Trails

Given the abundance of lEVs produced from the U-87 MG bioreactor cultures, we attempted to micropattern migrasome-mimetic trails mimicking those seen in several recent publications[28, 56, 57]. The U-87 MG lEVs used throughout this study were isolated using an established protocol for migrasome isolation from serum involving density gradients[31], but further characterization is required to say with confidence that these are mostly migrasomes rather than other lEV subtypes. Also, as this type of experiment was the first of its kind, the geometry and concentration of the the deposited EV trails were simplified in order to ascertain whether LEVA could be used to produce scalable platforms for studying the effects or internalization kinetics of surface-bound EVs on deposited cells. Following LEVA patterning, U-87 MG cell behavior was recorded using time-lapse epifluorescence microscopy. As shown in Fig 5 and in Supplementary Video 5, single U-87 MG cells clearly attached and migrated along the engineered regions that were treated with lEVs, exhibiting highly 1D migration behavior along the patterned regions. Supplementary Video 6 shows a similarly engineered but FBS-treated surface with less attachment and migration.

**Fig. 5.**
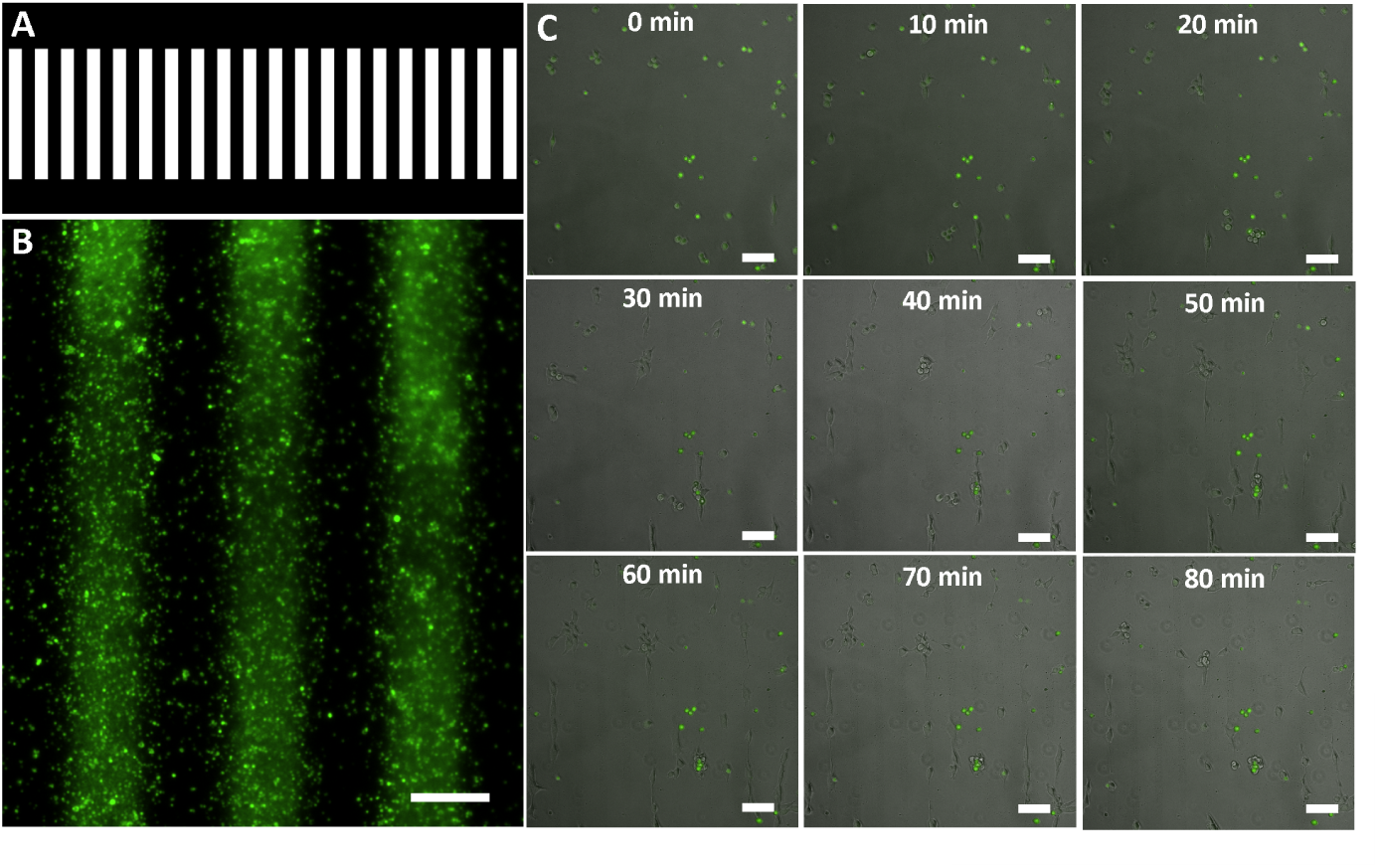
U-87 MG glioblastoma cell migration on migrasome-mimetic trails. (A) Template of 20 *µ*m wide microtracks used for engineering surface of cover slip bottom 96-well plate and (B) TIRFM image of resulting migrasome-mimetic trails made of U-87 lEVs (scale bar = 20 *µ*m). (C) Time-lapse images of U-87 MG cell migration on migrasome-mimetic trails (scale bar = 60 *µ*m).

### 1.6 OMV-mediated Neutrophil Swarming

To further display the relevance of LEVA for biomedical applications, bacterial OMVs were used as a stimulant for human peripheral blood-derived neutrophils. For validation of E. coli OMV size distribution and conformation, NTA and negative stain TEM were utilized as shown in Fig 6A. Then, to identify whether OMVs produced morphological differences in neutrophils in suspension, identified as a state of activation, OMVs and neutrophils were allowed to incubate together for 30 minutes before imaging using inverted epifluorescence microscopy. In Fig 6B phase-contrast images of native neutrophils (1), neutrophils with a WGA dye-only control (2), and neutrophils with WGA stained OMVs (3), demonstrate that OMV-exposed neutrophils underwent significantly more morphological changes. This morphological change is shown in the bar graph, where nearly 80% of neutrophils in this condition are activated.

**Fig. 6.**
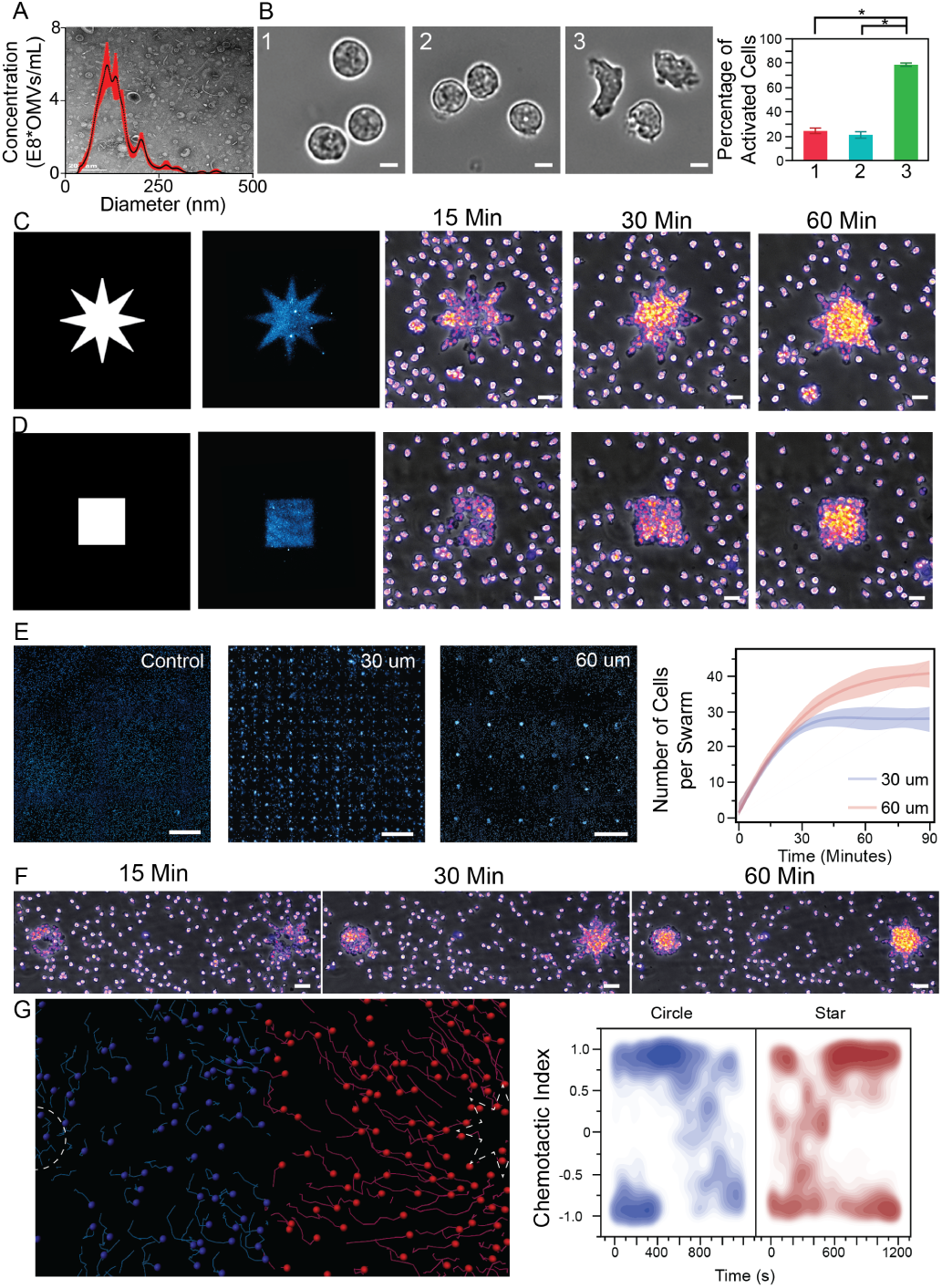
Study of OMV-mediated Neturophil Swarming. (A) NTA size distribution and TEM image of E. coli OMVs. (B) Phase contrast images of [1] native neutrophils, [2] neutrophils with WGA dye, and [3] neutrophils with WGA-labeled OMVs (scale bars = 5 *µ*m) with percentage activation of each condition on the right (* denotes p *<* 0.0001 Tukey’s HSD, n = 3 0.45mm^2^ FOV). (C-D) (left to right) Patterning template of star (above) and square (below) (both patterns are approximately 2830 *µ*m^2^), OMVs filled into the pattern, and time-lapse images of OMV-mediated neutrophil swarm at 15, 30 and 60 minutes, respectively (phase contrast and cell nucleus is the glow, scale bars = 10 *µ*m). (E) (left to right) Endpoint (90 minutes) wide FOV (2500 *µ*m x 2500 *µ*m) of Control, 30 *µ*m diameter OMV pattern neutrophil swarms (17×17), and 60 *µ*m OMV diameter pattern neutrophil swarms (6×6) (nucleus is cyan hot and scale bars = 400 *µ*m) and neutrophil accumulation over time on 30 *µ*m and 60 *µ*m OMV patterns (n=25 swarms). (F) Circle and star OMV pattern with 400 *µ*m center-to-center spacing (both patterns are approximately 2830 *µ*m^2^) time-lapse images of the OMV-mediated neutrophil swarming at 15, 30, and 60 minutes (phase contrast and cell nucleus is the glow and scale bars = 30 *µ*m). (G) Neutrophil migratory tracks after 20 minutes of swarming where the tracks are displayed either on the star half of the FOV (red) or circle half of the FOV (blue) and the neutrophil’s chemotactic index overtime either towards the circle OMV pattern (blue) or star OMV pattern (red).

As neutrophils are activated by OMVs, it was of great interest to determine whether surface-bound OMVs could initiate the neutrophil swarming cascade for the first time. Fig 6C-D displays the initial template utilized by LEVA, where an 8-point star is displayed in Fig 6C and a square in Fig 6D (surface area = 2830 *µ*m^2^ for all OMV patterns), the resulting OMV patterns imaged through TIRFM, and a 15, 30, and 60 minute time-point image of the OMV-mediated neutrophil swarms. A time-lapse video over the course of two-hours can be seen for the star and square OMV-mediated swarm in Supplementary Videos 7-8.

LEVA’s capabilities include the ability to micropattern EVs across large surface areas, and this is displayed by the large arrays of OMV-mediated neutrophil swarms in Fig 6E, where 2500 *µ*m x 2500 *µ*m fields of view (FOV) of an FBS-only control 17×17 30 *µ*m diameter 150 *µ*m center-to-center spacing pattern (far left), 17×17 array of the same pattern OMV induced neutrophil swarms, and 6×6 60 *µ*m diameter 400 *µ*m center-to-center spacing OMV induced neutrophil swarms. In these images, there are roughly 15 more neutrophils per swarm present in the 60 *µ*m diameter pattern compared to the 30 *µ*m diameter pattern after 90 minutes. A video of 30 *µ*m circle OMV-mediated neutrophil swarms can be seen in Supplementary Video 9.

To further the versatility of LEVA, multiple different geometric templates were utilized within the same microdomain. This can be seen in Fig 6F, where 15, 30, and 60 minute time-point images of a circle (left) and an 8-point star (right) are 400 *µ*m apart, with both patterns made with an equal area of 2830 *µ*m^2^. A time-lapse video over the course of two-hours can be seen in Supplementary Video 10. Interestingly, despite being the same surface area, it was observed that the cells’ migration differed when these two patterns were present in close proximity, as observed through the cells migratory tracks in Fig 6G, where the red spots represent cells which are on the star’s half of the FOV and the blue spots represent cells which are on the circle’s half. A video of the tracks can be seen in Supplementary Video 11, where the same colors are used as Fig 6G.

Interestingly, within the blue tracks there are a large amount which appear to be turning around shortly after swarming begins. This change in migratory behavior can be measured through the chemotactic index. Chemotactic index is a measurement of the neutrophils directionality, where a positive value indicates the cell is traversing towards the described OMV pattern, and a negative value indicates the cell is traversing away. The chemotactic index towards the circular OMV pattern (blue on the heatmap) demonstrates that after 600 seconds, there is a clear change in the directionality from moving towards the circular pattern to moving towards the star OMV pattern, whereas the chemotactic index towards the star (red on the heatmap) increases from 0-600 seconds and then has substantially larger amounts of cells migrating towards the OMV pattern. When the chemotactic index of a star-circle swarm is compared to a circle-circle swarm, as shown in Supplementary Video 12, it can be seen in the circle-circle swarm that there is no noticeable shift in directionality over time, which are apparent in the circle-star swarms’ chemotactic index.

## 2 Discussion

The reported results clearly establish the basic principles and limitations of the LEVA technique and demonstrate a small sample of the range of novel applications to which it can be applied. Single EV characterization by digital titration and fluorescence colocalization, single cell migration on migrasome or migrasome-mimetic trails, and OMV-mediated neutrophil swarming are only a few of the many EV and nanoparticle-related applications which are enabled by LEVA, indicating the potential value that this platform technology could provide to the EV field is limitless as EV researchers have begun to appreciate the role of ECM- and surface-bound EVs in human health and disease.

While the COMSOL simulation results rely on several assumptions to approximate the EV properties and underlying LEVA kinetics, the results are promising. This initial study was limited to only large and sEVs from a U-87 MG CELLine adherent bioreactor culture, but the similarities in trends from the simulations and time-lapse TIRF imaging indicate that further efforts to expand this model could be useful for other types of particles. In the future several other factors can be included in this approach to kinetics characterization, such as lipoprotein contamination, changes in EV corona due to processing or storage conditions, or the use of different buffers and surface materials[58–61].

Without question, one of the most valuable, but technically complex applications enabled by LEVA is the mimicking of migrasomes or other matrix- and surface-bound EVs followed by the culture of deposited cells. The scope of the results reported in this study are largely for proof-of-concept and limited to micropatterning of GBM lEVs and subsequent attachment and migration of GBM cells. However, progress towards ideal biomimicry of ECM-bound EVs, including migrasomes, will require significant optimization and tuning of the engineered surfaces for specific microenvironments. For example, in the case of migrasomes, this will include careful study of the EV-bound retraction fibers that are often found attached to the migrasomes’ pomegranate-like structures. In the future, the production and isolation of migrasomes, as well as their deposition will be improved to ensure that their morphology reflects their native state as closely as possible. These surface-bound EVs can then be compared to other engineered microenvironmental stimuli to compare their relative influence on cell behavior and function compared to other stimuli. Finally, in addition to engineering scalable, bioimimetic EV microenvironments, we anticipate that LEVA will improve our fundamental understanding of the internalization and signaling migrasomes and ECM-bound EVs, encouraging the development of relevant therapies therapies from applications ranging from cancer to wound healing based on these findings.

OMVs have been shown to elicit a stronger response from antigen presenting cells[62], but little is known about the change in neutrophil response when presented OMVs as compared to live bacteria[45] or antigen-coated microspheres[63]. Furthermore, this platform permits the multifaceted study of other immune cell-OMV interactions which are yet to be explored. For example, there have been studies aiming to understand the neutrophil-biofilm coexistence[64] and biofilms are understood to be continually releasing OMVs[65], therefore LEVA enables this relationship between biofilms, OMVs, and neutrophils (or other immune cells) to be better understood. Additionally, super-resolution microscopy of neutrophil swarms has been shown to identify the changes of organelles within the neutrophil crowd[66], and the LEVA platform will likely enhance these studies. LEVA is not only versatile in the type of EVs that can be used with the reported version of the surface patterning technique, but also the underlying surfaces themselves. For instance, during the initial LEVA steps when the surface is coated with PLL, it is possible that ECM proteins of interest could be used independently, or in combination with PLL to mimic different microenvironments. While the exact chemical location of photocleavage produced by the UV illumination is still unclear, it is likely that some, if not all of this initial layer remains on the surface and could be changed to mimic different microenvironments or modify the interactions with adsorbed EVs. In fact, any initial layer which allows subsequent functionalization with PEG by targeting amines should be possible with slight modifications to the reported approach. While not reported in this study, it is also possible to cleave any residual PEG following micropatterning, in a manner similar to sacrificial layer removal in photolithography by using flood exposure of the entire surface with UV illumination. However, this optional step requires optimization and careful consideration of the effects that UV exposure could have on the captured EVs.

Furthermore, in this study we largely performed patterning on glass cover slips to enable simple TIRFM imaging as required for continual validation in this type of study. However, based on our experience, as long as the general surface chemistry approach is the same and similar UV illumination is possible, LEVA could be applied to many other types of surfaces with similar resolution, tunability, and throughput. While this may limit TIRFM or other single EV-resolution imaging techniques, once the EV micropatterning is validated, a wide range of other functional surfaces could be patterned using LEVA. These could include engineered topographical cues and other 2D and 3D micro- and nanostructures, polymer thin films, and other surfaces to create multifunctional platforms and devices. In these cases electron microscopy may be required to validated LEVA, but once developed, these more advanced surfaces could also be incorporated into microfluidic and other bioMEMS devices to incorporate dynamic fluidic conditions for both the EVs and cells of interest. Finally, it is important to note that LEVA, to some degree, relies on a the researcher having basic knowledge of proper EV production, isolation, labeling, and characterization, which have been discussed at length in the MISEV guidelines[6] but are particularly important given the nonspecific nature of LEVA’s EV capture. Future studies which use LEVA to claim any functional effects of matrix- or surface-bound EVs should first clearly describe the EV isolation, characterization, and labeling methods used, as soluble protein or free dye contaminants will rapidly bind to the engineered surfaces and confound the actual effects of the EVs on the cells of interest. These issues have plagued research involving EVs in fluids and should be avoided as much as possible, enabling the responsible growth of this field.

## 3 Conclusions

In this study, we introduce a powerful new platform technology called Light-induced Extracellular Vesicle Adsorption (LEVA), enabling the efficient micropatterning EVs onto surfaces in a scalable, versatile, and high-resolution manner. LEVA should rapidly advance the study of matrix- and surface bound EVs from diverse sources and provide a clearer understanding of their functional role in cell-to-cell communication. The development and optimization of LEVA was enabled by the production of vast amounts of EVs from bioreactors and validated with commercially available GFP-EV standards and E. coli OMVs. While the underlying adsorption kinetics of EVs are still not fully understood, expanding on the reported simulation and time-lapse microscopy studies will enable more efficient patterning, and has already clearly established LEVA’s dependence on EV size. While this study focused on the basic validation of the LEVA process and demonstrating three novel applications, it is evident that LEVA will provide a platform for limitless other applications in the hands of capable EV researchers. Additionally, researchers aiming to implement LEVA must take great care to ensure responsible EV isolation and labeling given its nonspecific nature. LEVA should disrupt many current technical challenges in the study of matrix- and surface-bound EVs, whether for biomimicry or controlling cell behavior on biomaterials, and when used appropriately should improve rigor and reproducibility in this rapidly growing field.

## 4 Methods

### 4.1 Light-induced EV Adsorption

LEVA was developed by repurposing the previously reported Light-induced molecular adsorption of protein (LIMAP) technique [67], with careful considerations and adaptations needed for the EV field including EV production, isolation, characterization, labeling, and application-relevant patterning. First, glass coverslips were cleaned with ethanol and then deionized (DI) water via sonication for 3 min. The surface of the coverslip was treated with oxygen plasma for 1 min to activate the surface. A small drop of 0.01 % (w/v) poly-L-lysine (PLL) was placed onto parafilm, on which the treated coverslip was then placed for an even distribution of the PLL through physisorption. After incubating the coverslip for 30 min at room temperature, the PLL-coated coverslip was rinsed with DI water and dried with nitrogen flow. Following the same method, 100 mg/mL of methoxy-poly(ethylene glycol)-succinimidyl valerate (mPEG-SVA) diluted in 0.1 M HEPES was covalently bound to the PLL-coated coverslip to create a non-biofouling surface through N-hydroxysuccinimide (NHS) chemistry. The coverslip was incubated at room temperature for 1 h before rinsing with DI water and drying with a nitrogen airflow. 4-benzoylbenzyl-trimethylammonium chloride (PLPP) was then added as a photoactivator prior to UV illumination in a manner dependent on the specific EV application (PLPP gel was diluted in 96% ethanol in most single exposure cases).

The passivated coverslips were photoetched using a DMD optical module (Primo, Alveole) mounted on an automated inverted microscope (Nikon Eclipse Ti). Briefly, grayscale images were translated into high resolution UV light, allowing for a maskless illumination of different UV intensities correlating to the corresponding grayscale values. For initial testing, once the PLPP-ethanol evaporated, a silicone spacer (3.5 mm × 3.5 mm, 64 wells) was placed on the PEG-functionalized coverslip. For most LEVA experiments, grayscale values ranging from 30 to 50 % and UV doses of 20-30 mJ/mm^2^ were used. After the UV illumination, the photoetched coverslip was washed under a stream of DI water and dried by nitrogen flow. For most TIRFM imagine experiments, a 64-well ProPlate microarray system (Grace Bio Labs) was placed gently on the photoetched coverslip. The assembled array was secured by self-cut Delrin snap clips (Grace Bio Labs) to avoid leakage or potential contamination. The photoetched coverslip was rehydrated in PBS for 5 min and the EVs were allowed to adsorb for up to 10 min following 10 washes with PBS.

### 4.2 LEVA Simulations

The LEVA simulations were conducted using a COMSOL Multiphysics version 6.0 license provided by the Ohio State Supercomputer Center at The Ohio State University, Columbus, USA. We utilized the time-dependent Lagrangian Particle Tracing for fluid flow module to gain insights into the dynamics of EVs on the engineered glass surface. The glass surface measured 80 *µ*m x 80 *µ*m and featured nine equidistant 10 *µ*m patterns. These patterns were made positively charged, while the rest of the surface remained neutral and non-reactive. The EVs were considered as suspended particles in a PBS solvent with physical properties including a density of 1.13 mg/mL and a surface tension similar to PBS. Two types of samples were analyzed: sEVs with a mean size of 100 nm and a standard deviation of 50, and larger EVs with a mean size of 135 nm and a standard deviation of 45. The zeta potential of the EV particles was found to be negative, allowing them to bind with the positively charged patterns on the glass surface.

The simulations incorporated Lennard-Jones particle interactions coupled with Newtonian laws of motion for particle propagation. The initial boundary condition was set at time 0 sec, with all particles grid mesh-based release onto the entire surface. The walls were treated as diffusion scattering to ensure varied angles of reflection, providing a closer approximation to reality. The influence of gravity was neglected, as Brownian motion dominated the nanoscale particle movements. For the Finite Particle Tracing (FPT) simulations, we used a physics-controlled extremely fine mesh. The time-dependent equations were solved using 20 vertex elements, 560 boundary elements, and a total of 28,834 domain elements. These computations were distributed across 4 nodes with 112 cores and took 13 hours to calculate trajectories of 2.78E8 particle concentration per mL with an output time of up to 60 seconds, in steps of 0.0001. Post-processing involved calculating the average surface density of particles and the surface integral of velocity magnitude. Particle trajectories at various time intervals were collected and analyzed to determine particle density at the binding sites.

### 4.3 Commercial EV Standards

Commercially available GFP-EV standards were used for the final validation of LEVA. The lyophilized GFP-EV standards were reconstituted by carefully following the specific manufacturer’s instructions. Vesi-Ref EVs (Vesiculab) and fluorescent exosome standards (SAE0193, Sigma) were resuspended in 100 ul of ultrapure water, avoiding bubbles. Vesi-Ref EVs were then vortexed for 60 seconds, briefly centrifuged, and briefly mixed again via pipetting. Concentrations of both commercial EV standards were then normalized by NTA measurement prior to patterning.

### 4.4 Bioreactor Production of Cell Line EVs

CELLine AD 1000 bioreactors (Wheaton) were used for EV production following an established adaptation and EV collection and isolation protocol [54, 55, 68]. Briefly, bioreactors were adapted from complete media with 10 % fetal bovine serum (FBS) supplementation, then adapted to CDM-HD (Fiber-cell). EVs were collected in around 15 ml of conditioned media three times per week from the cell chamber, and the media chamber was refreshed once per week.

### 4.5 EV Isolation and Labeling

Cells and large debris were first removed using differential centrifugation via a 200 × g for 5 min followed by a 2000 × g for 10 min. lEVs and sEVs were then isolated using a combination of differential ultracentrifugation and either density gradient ultracentrifugation or size exclusion chromatography. Crude lEVs were isolated by ultracentrifugation (Beckman XP-100, Ti 70 rotor) at 20,000 × g for 30 min, and the resulting supernatant was then ultracentrifuged (Beckman XP-100, Ti 70 rotor) at 100,000 × g for 60 min to isolate crude sEVs. The lEV pellets were further fractionated using an iodixanol (OptiPrep, D1556-250ML) density gradient containing 500 *µ*L layers of 30 %, 25 %, 19 %, 15 %, 12 %, 10 %, 8 %, and 5 % with the lEVs contained in the top layer. Following ultracentrifugation (Beckman XP-100, SW 28 Ti rotor) at 150,000 × g for 4 h, lEVs were isolated from seventh fraction 7 by washing twice with PBS by centrifugation at 10,000 xg for 30 min each wash. The sEVs were isolated by pooling fractions 1-4 following size exclusion chromatography using a 35 nm qEV Original column coupled with an Automated Fraction Collector (AFC, Izon). All EVs were characterized using basic NTA (Nanosight NS300). Given previously reported successes in labeling migrasomes with WGA[69], lEVs and sEVs were incubated with 1*µ*g/ml wheat germ agglutinin (WGA kit, W7024, Thermo) for 10 min at room temperature. Excess dye was removed via ultrafiltration using 100 kDa filters (Vivaspin 500, Sartorious) at 10,000 × g. The solution was fully washed with PBS three times following the same procedure, including a dye only control.

### 4.6 Production of GFP containing OMVs from E. coli

The pET/EGFP-strep-his plasmid used to express the green fluorescent protein (GFP) containing outer membrane vesicles (OMVs) was acquired from the Wood Lab strain collection (Ohio State University, OH, USA). Enhanced green fluorescent protein (EGFP) has a monomerizing single A206K mutation [70]. pET/EGFP-strep-his was miniprepped according to manufacturer’s instructions (Qiagen) and transformed into competent Escherichia coli BLR (DE3) cells. The primary culture (10 mL Luria Broth media supplemented with 100 ug/mL ampicillin) was inoculated with a single transformed colony and grown at 37°C and 225 rpm for 18 h. The 10 mL primary culture was used to inoculate one liter of Terrific Broth (TB) media (1 % v/v inoculum) supplemented with 100 ug/mL ampicillin in a 2.5 L Thompson Ultra Yield™ flask (Oceanside, CA, USA). The culture was grown to an OD600 of 3 ± 0.5 at 37°C and 275 rpm and then induced with 0.5 mM (final concentration) of isopropyl *β*-D-1-thiogalactopyranoside (IPTG). The culture was left shaking for 16 hours at 16°C and then harvested at 8,000 rcf for 20 minutes. The supernatant containing the OMVs (conditioned media) was collected and frozen at -20°C until ready for use.

To begin concentrating the OMVs, the conditioned media was thawed at 4°C and 0.2 *µ*m sterile filtered using a vacuum flask (Corning, Cat#430769). The supernatant was then 10-times enriched through tangential flow filtration, which used a polysulfone hollow fiber cartridges (Molecular weight cutoff: 300kDa, Repligen, Cat#D02-S500-05-N) as the filter. The enriched supernatant was then further purified through size exclusion chromatography in the same manner as bioreactor-produced U-87 MG sEVs (qEV Original, Izon). Importantly, initial resolution testing of LEVA patterning with OMVs was performed by WGA-488 labeling, whereas neutrophil swarming experiments were performed with GFP-OMVs to reduce any potential confounding issues caused by the WGA dye. A schematic of all of the EV production isolation protocols can be seen in Supplementary Figure 2.

### 4.7 Human Neutrophil Isolation and Labeling

The morning of an experiment, human blood was collected in K2-Ethylenediaminetetraacetic acid (EDTA), purple top, tubes (BD Vacutainer, Fisher Scientific) according to protocol #2018H0268 approved by the Biomedical Sciences Committee Institutional Review Board (IRB) at The Ohio State University. The fresh blood was added to a 10 mM sodium bicarbonate, 0.1 mM EDTA, and 150mM ammonium chloride solution of pH 7.4 at a 1:20 ratio for 5 minutes to lyse the red blood cells. The solution was then centrifuged at 350 x g to isolate the white blood cells and remove the red blood cell debris. Neutrophils were retained in solution through negative immuno-magnetic separation through the EasySep™ Human Neutrophil Isolation Kit (STEMCELL Technologies, Vancouver, Canada). Neutrophils were incubated in Iscove’s Modified Dulbecco’s Medium (IMDM, ThermoFisher), and stained with 20*µ*g/mL Hoechst 33342 (Thermofisher) for 10 minutes. The neutrophils were then washed by adding phosphate buffer solution (PBS) to 5-times the original cell volume and centrifuged at 1000 RPM. The final neutrophil suspension was at a concentration of 1.2 x 10^6^ and contained IMDM, 20% fetal bovine serum (FBS) (Gibco, Thermofisher), and 1% penicillin-streptomycin (PS) (Gibco, Thermofisher).

### 4.8 EV Characterization

EVs were characterized primarily using NTA and TEM (Tecnai). For NTA, 5 x 30 second videos were recorded (acquisition: screen gain 1, camera level 15) at low flow rates and analyzed using the v3.3 software (screen gain 10, detection threshold 5). For negative staining TEM, on a strip of parafilm, 20 *µ*L droplets of water for injection (WFI) and two 20 *µ*L droplets of negative stain (UranyLess EM stain, Electron Microscopy Sciences) were placed. The TEM grid was subjected to plasma treatment for 1 min. 10 *µ*L of the pooled EV solution was placed onto the treated surface of the grid then was incubated for 1 min and blotted with filter paper to remove excess. The grid was then submerged into the WFI droplet and blotted dry using filter paper, and this process was repeated with the second WFI droplet. The grids were stained via submersion into the negative stain, blotted, and then submersion into the second droplet of the negative stain. The grid was incubated in the stain for approximately 22 seconds and excess solution was again wicked away using filter paper. Grids were stored overnight in a grid box to ensure thorough drying. TEM imaging was performed using a Tecnai TF-20 microscope (FEI Company, Hillsboro, OR) operating at 200 kV. Images of fluorescently labeled siEVPs were obtained by TIRFM (Nikon Eclipse Ti) with a 100× oil immersion lens.

## Supplementary information

The Supplementary Information contains additional figures including dye-only controls of all LEVA patterns, videos of both COMSOL Multiphysics simulations and time-lapse TIRFM demonstrating EV binding for lEVs and sEVs, videos of cell migration on migrasome-mimetic trails, as well as videos of neutrophil swarming and videos of neutrophil swarming analyses.

## Acknowledgments

This work was supported by The Ohio State University Center for Cancer Engineering-Curing Cancer Through Research in Engineering Sciences, the National Institutes of Health grants: K99EB033857, UG3/UH3TR002884, U18TR003807, and the LEGACY Postdoctoral Scholars Program at Ohio State. SM would like to thank Professor Angela Brown at Lehigh University for encouraging her interest in OMV research. We would also like to thank our healthy donors whose blood donations enabled the neutrophil studies.

## Declarations

The authors have no conflicts of interest to declare.

## References

[1] Toyofuku, M., Nomura, N. & Eberl, L. Types and origins of bacterial membrane vesicles. Nature Reviews Microbiology 17 (1), 13–24 (2019).

[2] Yáñez-Mó, M., et al. Biological properties of extracellular vesicles and their physiological functions. Journal of extracellular vesicles 4 (1), 27066 (2015).

[3] Li, S.-R. et al. Tissue-derived extracellular vesicles in cancers and non-cancer diseases: Present and future. Journal of Extracellular Vesicles 10 (14), e12175 (2021).

[4] Couch, Y. et al. A brief history of nearly everything–the rise and rise of extracellular vesicles. Journal of extracellular vesicles 10 (14), e12144 (2021).

[5] Kalluri, R. & LeBleu, V. S. The biology, function, and biomedical applications of exosomes. Science 367 (6478), eaau6977 (2020).

[6] Welsh, J. A. et al. Minimal information for studies of extracellular vesicles (misev2023): From basic to advanced approaches. Journal of extracellular vesicles 13 (2), e12404 (2024).

[7] van Niel, G. et al. Challenges and directions in studying cell–cell communication by extracellular vesicles. Nature Reviews Molecular Cell Biology 23 (5), 369–382 (2022).

[8] Bordanaba-Florit, G., Royo, F., Kruglik, S. G. & Falcon-Perez, J. M. Using single-vesicle technologies to unravel the heterogeneity of extracellular vesicles. Nature Protocols 16 (7), 3163–3185 (2021).

[9] Hilton, S. H. & White, I. M. Advances in the analysis of single extracellular vesicles: A critical review. Sensors and actuators reports 3, 100052 (2021).

[10] Kwon, Y. & Park, J. Methods to analyze extracellular vesicles at single particle level. Micro and Nano Systems Letters 10 (1), 14 (2022).

[11] Lennon, K. M. et al. Single molecule characterization of individual extracellular vesicles from pancreatic cancer. Journal of Extracellular Vesicles 8 (1), 1685634 (2019).

[12] Jørgensen, M. M., Bæk, R. & Varming, K. Potentials and capabilities of the extracellular vesicle (ev) array. Journal of extracellular vesicles 4 (1), 26048 (2015).

[13] Saftics, A. et al. Single extracellular vesicle nanoscopy. Journal of Extracellular Vesicles 12 (7), 12346 (2023).

[14] Panagopoulou, M. S., Wark, A. W., Birch, D. J. & Gregory, C. D. Phenotypic analysis of extracellular vesicles: a review on the applications of fluorescence. Journal of extracellular vesicles 9 (1), 1710020 (2020).

[15] Zhang, J. et al. Engineering a tunable micropattern-array assay to sort single extracellular vesicles and particles to detect rna and protein in situ. Journal of Extracellular Vesicles 12 (11), 12369 (2023).

[16] Nguyen, L. T. et al. An immunogold single extracellular vesicular rna and protein (auserp) biochip to predict responses to immunotherapy in non-small cell lung cancer patients. Journal of Extracellular Vesicles 11 (9), e12258 (2022).

[17] McNamara, R. P. et al. Imaging of surface microdomains on individual extracellular vesicles in 3-d. Journal of Extracellular Vesicles 11 (3), e12191 (2022).

[18] Mizenko, R. R. et al. Tetraspanins are unevenly distributed across single extracellular vesicles and bias sensitivity to multiplexed cancer biomarkers. Journal of nanobiotechnology 19 (1), 250 (2021).

[19] Crescitelli, R., Lässer, C. & Lötvall, J. Isolation and characterization of extracellular vesicle subpopulations from tissues. Nature Protocols 16 (3), 1548–1580 (2021).

[20] Infanger, D. W., Lynch, M. E. & Fischbach, C. Engineered culture models for studies of tumor-microenvironment interactions. Annual review of biomedical engineering 15, 29–53 (2013).

[21] Hisey, C. L., Hearn, J. I., Hansford, D. J., Blenkiron, C. & Chamley, L. W. Micropatterned growth surface topography affects extracellular vesicle production. Colloids and Surfaces B: Biointerfaces 203, 111772 (2021).

[22] Huang, G. et al. Functional and biomimetic materials for engineering of the three-dimensional cell microenvironment. Chemical reviews 117 (20), 12764–12850 (2017).

[23] Sleeboom, J. J., Eslami Amirabadi, H., Nair, P., Sahlgren, C. M. & Den Toonder, J. M. Metastasis in context: modeling the tumor microenvironment with cancer-on-a-chip approaches. Disease models & mechanisms 11 (3), dmm033100 (2018).

[24] Debnath, K., Las Heras, K., Rivera, A., Lenzini, S. & Shin, J.-W. Extracellular vesicle–matrix interactions. Nature Reviews Materials 8 (6), 390–402 (2023).

[25] Rilla, K. et al. Extracellular vesicles are integral and functional components of the extracellular matrix. Matrix Biology 75, 201–219 (2019).

[26] Rackov, G. et al. Vesicle-mediated control of cell function: The role of extracellular matrix and microenvironment. Frontiers in physiology 9, 651 (2018).

[27] Lewin, S., Hunt, S. & Lambert, D. W. Extracellular vesicles and the extracellular matrix: a new paradigm or old news? Biochemical Society Transactions 48 (5), 2335–2345 (2020).

[28] Ma, L. et al. Discovery of the migrasome, an organelle mediating release of cytoplasmic contents during cell migration. Cell research 25 (1), 24–38 (2015).

[29] Wu, D. et al. Pairing of integrins with ecm proteins determines migrasome formation. Cell Research 27 (11), 1397–1400 (2017).

[30] Sung, B. H., Parent, C. A. & Weaver, A. M. Extracellular vesicles: Critical players during cell migration. Developmental cell 56 (13), 1861–1874 (2021).

[31] Zhao, X. et al. Identification of markers for migrasome detection. Cell discovery 5 (1), 27 (2019).

[32] Shapiro, I. M., Landis, W. J. & Risbud, M. V. Matrix vesicles: are they anchored exosomes? Bone 79, 29–36 (2015).

[33] Lindenbergh, M. F. & Stoorvogel, W. Antigen presentation by extracellular vesicles from professional antigen-presenting cells. Annual Review of Immunology 36, 435–459 (2018).

[34] Lu, M. et al. The role of extracellular vesicles in the pathogenesis and treatment of autoimmune disorders. Frontiers in Immunology 12, 566299 (2021).

[35] Srivastava, A., Rathore, S., Munshi, A. & Ramesh, R. Extracellular vesicles in oncology: from immune suppression to immunotherapy. The AAPS Journal 23, 1–15 (2021).

[36] Czernek, L. & Düchler, M. Functions of cancer-derived extracellular vesicles in immunosuppression. Archivum immunologiae et therapiae 24 Light-induced Extracellular Vesicle Adsorption experimentalis 65, 311–323 (2017).

[37] Rossaint, J. et al. Directed transport of neutrophil-derived extracellular vesicles enables platelet-mediated innate immune response. Nature communications 7 (1), 13464 (2016).

[38] Chen, Z., Larregina, A. T. & Morelli, A. E. Impact of extracellular vesicles on innate immunity. Current opinion in organ transplantation 24 (6), 670 (2019).

[39] Zhou, X. et al. The function and clinical application of extracellular vesicles in innate immune regulation. Cellular & molecular immunology 17 (4), 323–334 (2020).

[40] Díaz-Godínez, C., Ŕıos-Valencia, D. G., Garćıa-Aguirre, S., Martínez-Calvillo, S. & Carrero, J. C. Immunomodulatory effect of extracellular vesicles from entamoeba histolytica trophozoites: Regulation of nets and respiratory burst during confrontation with human neutrophils. Frontiers in Cellular and Infection Microbiology 1611 (2022).

[41] Lanyu, Z. & Feilong, H. Emerging role of extracellular vesicles in lung injury and inflammation. biomedicine & pharmacotherapy 113, 108748 (2019).

[42] Lämmermann, T., et al. Neutrophil swarms require ltb4 and integrins at sites of cell death in vivo. Nature 498 (7454), 371–375 (2013).

[43] Kienle, K. & Lämmermann, T. Neutrophil swarming: an essential process of the neutrophil tissue response. Immunological reviews 273 (1), 76–93 (2016).

[44] Walters, N., Nguyen, L. T., Zhang, J., Shankaran, A. & Réategui, E. Extracellular vesicles as mediators of in vitro neutrophil swarming on a large-scale microparticle array. Lab on a Chip 19 (17), 2874–2884 (2019).

[45] JunǵaKim, J., et al. Large-scale patterning of living colloids for dynamic studies of neutrophil–microbe interactions. Lab on a Chip 18 (11), 1514– 1520 (2018).

[46] Walters, N. et al. Analyzing inter-leukocyte communication and migration in vitro: neutrophils play an essential role in monocyte activation during swarming. Frontiers in Immunology 12, 671546 (2021).

[47] Réategui, E., et al. Microscale arrays for the profiling of start and stop signals coordinating human-neutrophil swarming. Nature biomedical engineering 1 (7), 0094 (2017).

[48] Kaparakis-Liaskos, M. & Ferrero, R. L. Immune modulation by bacterial outer membrane vesicles. Nature Reviews Immunology 15 (6), 375–387 (2015).

[49] White, J. R. et al. The complex, bidirectional role of extracellular vesicles in infection. Biochemical Society Transactions 49 (2), 881–891 (2021).

[50] Cecil, J. D., Sirisaengtaksin, N., O’Brien-Simpson, N. M. & Krachler, A. M. Outer membrane vesicle-host cell interactions. Microbiology spectrum 7 (1), 7–1 (2019).

[51] Burgelman, M., Vandendriessche, C. & Vandenbroucke, R. E. Extracellular vesicles: A double-edged sword in sepsis. Pharmaceuticals 14 (8), 829 (2021).

[52] Buzas, E. I. The roles of extracellular vesicles in the immune system. Nature Reviews Immunology 23 (4), 236–250 (2023).

[53] Almeria, C., Kreß, S., Weber, V., Egger, D. & Kasper, C. Heterogeneity of mesenchymal stem cell-derived extracellular vesicles is highly impacted by the tissue/cell source and culture conditions. Cell & Bioscience 12 (1), 1–15 (2022).

[54] Hisey, C. L. et al. Investigating the consistency of extracellular vesicle production from breast cancer subtypes using celline adherent bioreactors. Journal of Extracellular Biology 1 (9), e60 (2022).

[55] Artuyants, A. et al. in Production of extracellular vesicles using a celline adherent bioreactor flask 183–192 (Springer, 2021).

[56] Dharan, R. et al. Tetraspanin 4 stabilizes membrane swellings and facilitates their maturation into migrasomes. Nature Communications 14 (1), 1037 (2023).

[57] Fan, C. et al. Cell migration orchestrates migrasome formation by shaping retraction fibers. Journal of Cell Biology 221 (4), e202109168 (2022).

[58] Buźas, E. I., Tóth, E. Á., Śodar, B. W. & Szabó-Taylor, K. É. Molecular interactions at the surface of extracellular vesicles, Vol. 40, 453–464 (Springer, 2018).

[59] Wolf, M. et al. A functional corona around extracellular vesicles enhances angiogenesis, skin regeneration and immunomodulation. Journal of extracellular vesicles 11 (4), e12207 (2022).

[60] Tóth, E. Á., et al. Formation of a protein corona on the surface of extracellular vesicles in blood plasma. Journal of extracellular vesicles 10 (11), e12140 (2021).

[61] Śodar, B. W., et al. Low-density lipoprotein mimics blood plasma-derived exosomes and microvesicles during isolation and detection. Scientific reports 6 (1), 24316 (2016).

[62] Prior, J. T. et al. Bacterial-derived outer membrane vesicles are potent adjuvants that drive humoral and cellular immune responses. Pharmaceutics 13 (2), 131 (2021).

[63] Walters, N. & Réategui, E. Bioparticle microarrays for chemotactic and molecular analysis of human neutrophil swarming in vitro. JoVE (Journal of Visualized Experiments) (156), e60544 (2020).

[64] Hirschfeld, J. Dynamic interactions of neutrophils and biofilms. Journal of oral microbiology 6 (1), 26102 (2014).

[65] Wang, W., Chanda, W. & Zhong, M. The relationship between biofilm and outer membrane vesicles: a novel therapy overview. FEMS microbiology letters 362 (15), fnv117 (2015).

[66] Glaser, K. M., et al. Arp2/3 and the pentose phosphate pathway regulate late phases of neutrophil swarming. iScience 27 (1) (2024).

[67] Strale, P.-O. et al. Multiprotein printing by light-induced molecular adsorption. Advanced Materials 28 (10), 2024–2029 (2016).

[68] Tasma, Z. et al. Production of extracellular vesicles from equine embryo-derived mesenchymal stromal cells. Reproduction 164 (4), 143–154 (2022).

[69] Chen, L., Ma, L. & Yu, L. Wga is a probe for migrasomes. Cell discovery 5 (1), 13 (2019).

[70] Zacharias, D. A., Violin, J. D., Newton, A. C. & Tsien, R. Y. Partitioning of lipid-modified monomeric gfps into membrane microdomains of live cells. Science 296 (5569), 913–916 (2002).

