## Supplementary Figure 1 for "Light-induced Extracellular Vesicle Adsorption"

**Contents:**

**Supplementary Figure 1.** TIRFM Images from Resolution Test with Small EV and WGA dye-only control ............ **S3**
**Supplementary Figure 2.** EV Production and Isolation Workflow ............................................................................ **S4**

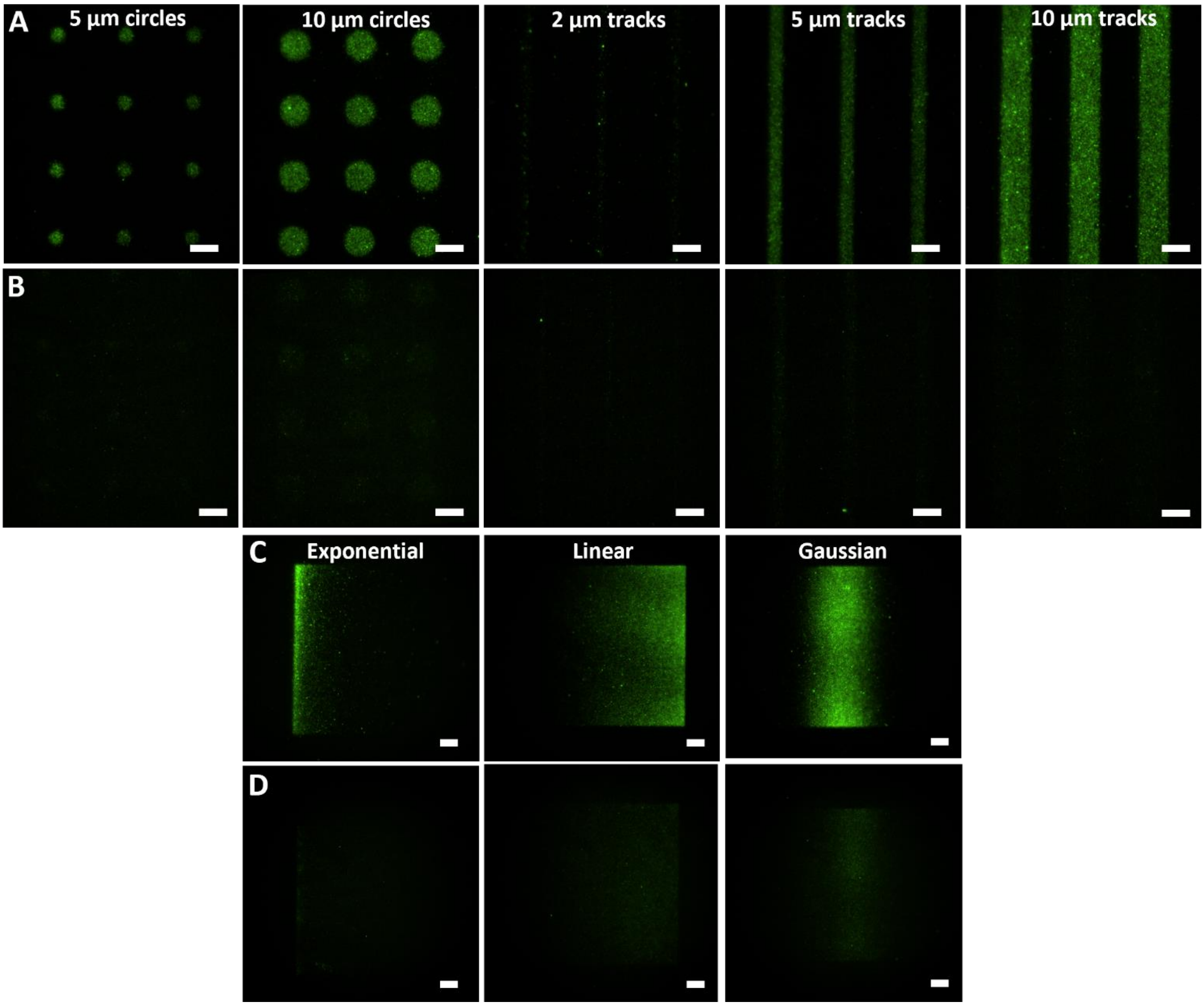


**Supplementary Figure 1.** LEVA resolution test comparing dye-only control. Small EVs and dye-only control were imaged using TIRFM with constant LUT values to demonstrate minimal background fluorescence. Resolution test results for 5 and 10 µm circle array template and 2, 5, and 10 µm microtrack array template shown for (A) WGA-labeled U-87 MG small EVs and (B) WGA dye-only control. Exponential, linear, and gaussian gradient LEVA tests for (C) WGA-labeled U-87 MG small EVs and (D) WGA dye-only control.

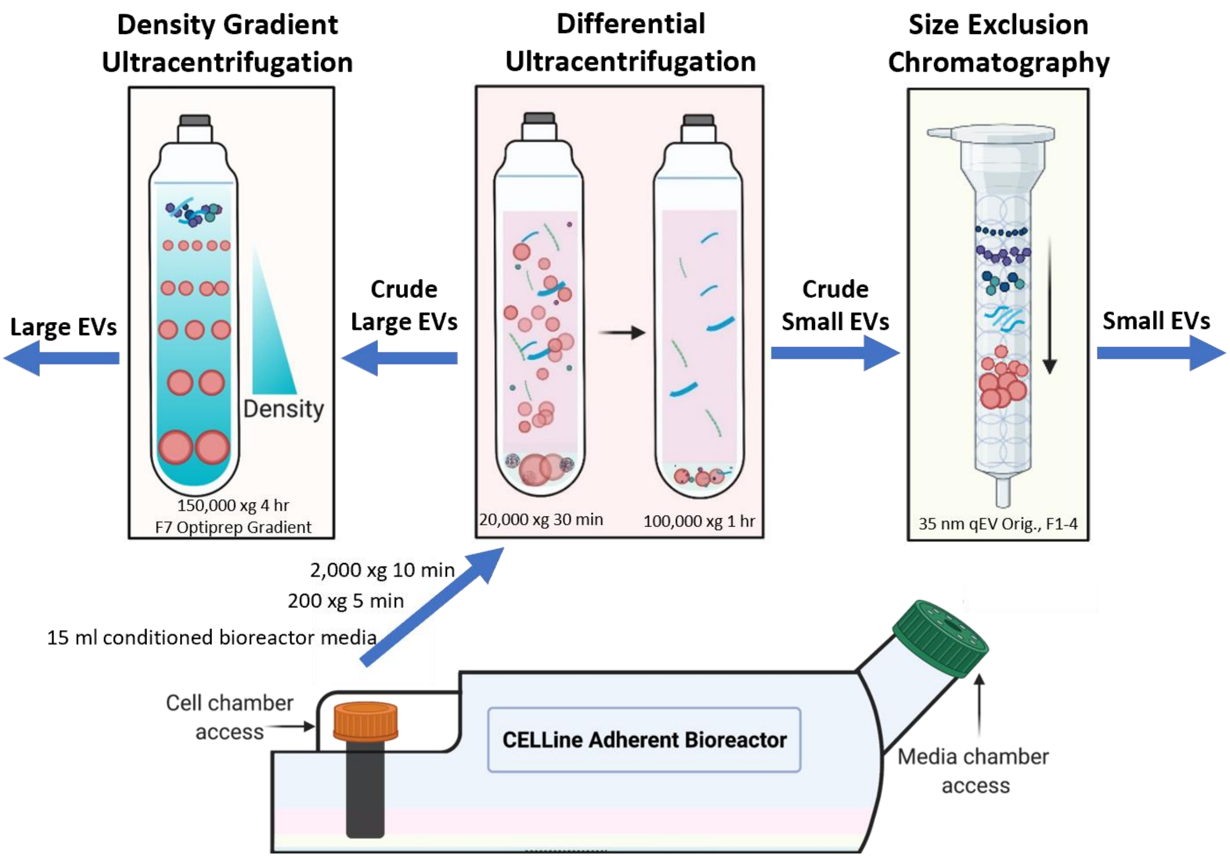


**Supplementary Figure 2:** EV Isolation Schematic for U-87 MG Large and Small EVs, showing collection of roughly 15 ml of conditioned media from CELLine Adherent Bioreactor cell chamber followed by centrifugation steps to remove cells and large debris, differential ultracentrifugation to isolate crude large and small EVs, and density gradient ultracentrifugation (DG) and size exclusion chromatography (SEC) to isolate large and small EVs, respectively, from other contaminants. The Optiprep DG used to isolate large EVs follows the "Isolation of migrasomes from serum samples" protocol established by Zhao *et al.* [31], and the SEC protocol uses a 35 nm qEV Original column with fractions 1-4 pooled, as detailed in Hisey *et al.* [54].
